# Spatially selective open loop control of magnetic microrobots for drug delivery

**DOI:** 10.1101/2023.03.31.535118

**Authors:** Nima Mirkhani, Michael G. Christiansen, Tinotenda Gwisai, Stefano Menghini, Simone Schuerle

## Abstract

Rotating magnetic fields (RMFs), when used to actuate biomedical microrobots for targeted delivery to tumors, have been shown to enable them to overcome physiological barriers and promote their accumulation and penetration into tissue. Nevertheless, directly applying a RMF to a deeply situated target site also leads to off-target actuation in surrounding healthy tissue. Here, we investigate an open-loop control strategy for delivering torque density to diffuse distributions of microrobots at focal points by combining RMFs with magnetostatic gating fields. Taking magnetotactic bacteria (MTB) as a model biohybrid microrobotic system for torque-based actuation, we first use simulation to elucidate off-target torque suppression and find that resolution is set by the relative magnitude of the magnetostatic field and RMF. We study focal torque delivery in vitro, observing off-target suppression of translational velocity of MTB, convection-driven accumulation of companion nanoparticles, and tumor spheroid colonization. We then design, construct, and validate a mouse-scale torque-focusing apparatus incorporating a permanent magnet array, three-phase RMF coils, and offset coils to maneuver the focal point. Our control scheme enables the advantages of torque-based actuation to be combined with spatial targeting, and could be broadly applied to other microrobotic designs for improved drug delivery.

**One-Sentence Summary:** Combining rotating magnetic fields with gating fields enables focused delivery of torque density to dispersed microrobots.

## INTRODUCTION

Delivering therapeutic payloads or immune-modulating living vectors to sites where they are needed remains a perennial challenge, especially in cancer therapy.^1^ The efficiency of delivery as a fraction of administered dose remains limited, even for carefully designed nanocarriers employing physiochemical targeting strategies.^2–5^ Similar hurdles have been encountered by self-propelled therapeutic bacteria, for which delivery efficiency is especially crucial because tolerable doses are low.^6–9^ Recently, the scope of investigation into biomedical microrobots suited for drug delivery has expanded remarkably.^10–12^ Microrobots designed to convert externally applied magnetic stimuli to locomotion, convection, or other modes of therapeutic activity are particularly useful for deep physiological targets, which magnetic fields can access unencumbered.^13–16^ Historically, magnetic field gradients that apply forces to magnetic materials introduced into the body have been envisioned for targeting,^17, 18^ an approach that is fundamentally limited to superficial targets in the absence of additional constraints.^19^ More recently, uniform fields that steer self-propelling microbots^11, 20–23^ or rotating fields that power motion through applied magnetic torques^24–29^ have offered compelling alternatives that are suited for deep tissue and scalable to patients (**Fig. 1A**). We previously showed how rotating magnetic fields (RMFs) can be applied to naturally magnetic bacteria, known as magnetotactic bacteria (MTB), to help them overcome physiological barriers to reach tumors. MTB can thus be seen both as biohybrid microrobots and potential vectors in bacterial cancer therapy.^30^

**Fig. 1.**
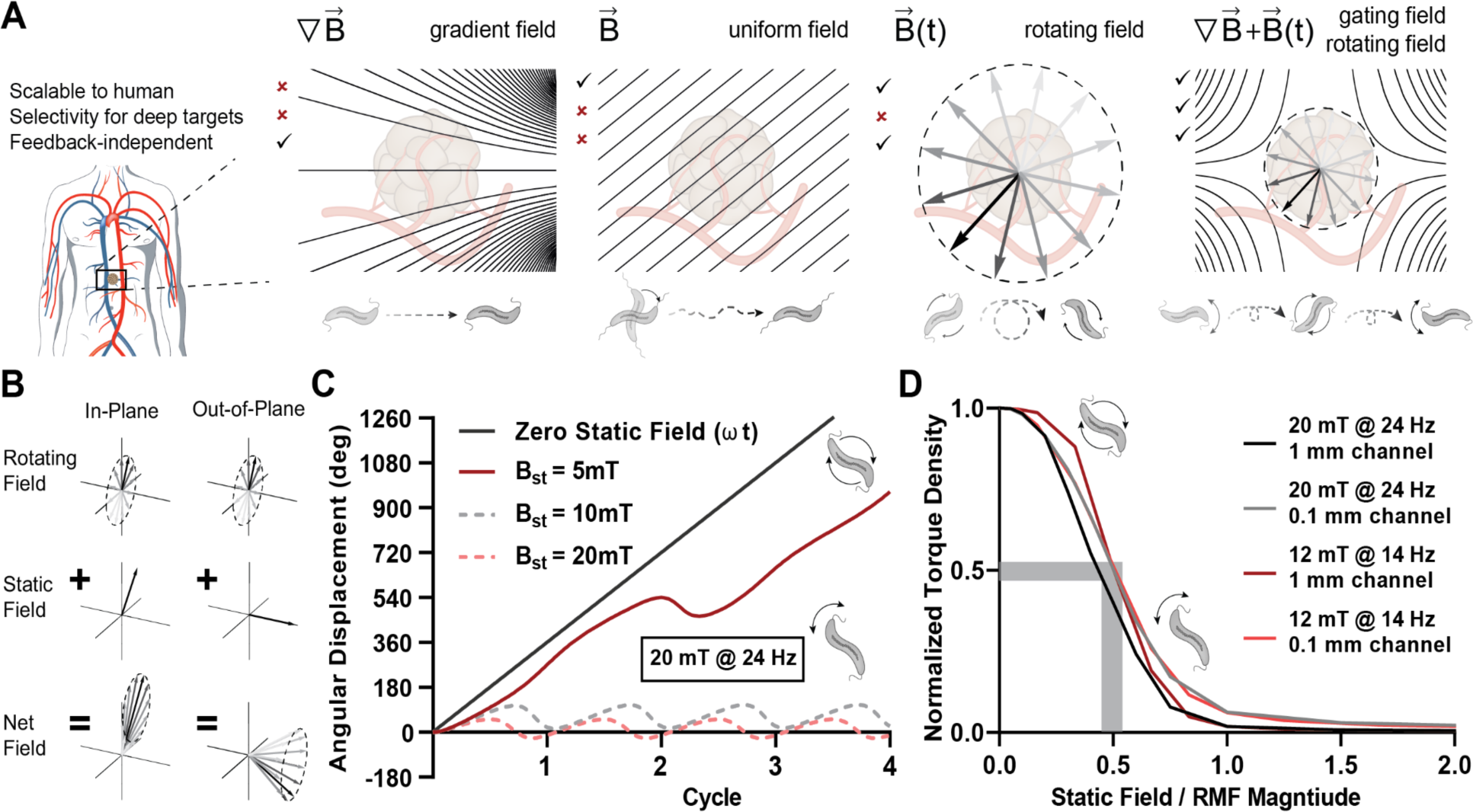
Concept of magnetic torque focusing by combining RMFs and magnetostatic gating fields. (**A**) Overview comparing the advantages and disadvantages of various forms of magnetic field application to control vascularly dispersed magnetic microrobots, here depicted as MTB, our model microrobot. Possible magnetic stimuli include gradient forces, uniform fields for steering, global RMFs, and RMFs with gating fields. (**B**) Vector superposition of magnetostatic fields and RMFs with a relative magnitude of 1.5 times for the special cases of a magnetostatic field vector in the plane of rotation (in-plane) and perpendicular to the plane of rotation (out-of-plane). (**C**) Angular displacement of simulated individual MTB microrobots exposed to different magnitudes of the static gating field with RMF of 20 mT and 24 Hz. (**D**) Simulated nondimensionalized torque density curves of bacteria under different geometric and actuation conditions. The curves nearly overlap, suggesting a generalized interpretation of off-target torque density suppression as determined by the relative magnitude of the magnetostatic field and RMF.

Given that RMFs, when applied to diffuse dispersions of synthetic or living microbots, are capable of enhancing extravasation and tissue penetration, it follows that a strategy to locally focus magnetic torque density is needed to reduce off target actuation. Ideally, this would utilize a magnetic stimulus that does not require detailed knowledge of the instantaneous distribution of microbots, but rather serves as a form of open loop control, selectively delivering torque density to microrobots within a target region as they circulate throughout the body (Fig. 1A). To accomplish this, a magnetostatic gating field that supplies a field free point or field free line can be introduced, which suppresses torque-based actuation outside these regions.^30–36^ This concept is adapted from magnetic particle imaging (MPI), where acquisition of inductive signals from diffuse magnetic nanomaterial is locally confined to image voxels by a similar selection field.^37–39^ More recently, research has also indicated the possibility of using magnetostatic fields superimposed with high frequency (100s of kHz) alternating magnetic fields to spatially restrict hysteretic heat dissipation of magnetic nanomaterials.^36, 40–44^

Here, we make use of selection fields as an integral component of a spatially targeted open loop control strategy for diffuse living microrobots via torque-based actuation. Using an in vitro platform based on a commercial multi-axis electromagnet setup in combination with an array of small permanent magnets, we demonstrate the possibility for localized actuation of MTB within a targeted region. We show how this targeted actuation influences translational velocity of the MTB, penetration of companion nanoparticles (NPs) into collagen matrices, and RMF-enhanced colonization of tumor spheroids. Encouraged by these results, we designed and constructed a torque focusing setup for small rodents, consisting of an external array of ferrite permanent magnets arranged as modified “magic sphere”, an internal three-phase RMF generator, and a pair of DC coils to adjust the position of the field free point. The position of the field free point along the longitudinal axis of the animal is maneuvered by a positioning track, while the position of the field free point within the transverse plane is adjusted electronically by currents within the DC coils. This device is validated through characterization of the net fields experienced in the working volume and through a magnetic mixing experiment that verified spatially selective convection. This strategy for torque focusing represents a further advantage of RMF-based actuation of biomedical magnetic microrobots, and provides a means to increase spatial selectivity in the effort to address longstanding challenges in targeted drug delivery.

## RESULTS

### Combining rotating and magnetostatic gating fields focuses torque application

Magnetic microrobots introduced intravenously typically are immediately dispersed throughout the body by the circulatory system, and magnetic stimuli can then be applied as part of a control strategy to influence their distribution. Although many approaches for magnetic control are possible, an ideal control scheme should not require knowledge of the instantaneous distribution of the microrobots, should scale well to form factors compatible with human patients, and should be able to specifically target remote points. Combining rotating magnetic fields with magnetostatic gating fields for selective torque-based actuation^30, 32, 36^ represents a way to simultaneously satisfy these requirements when applied to diffuse distributions of microrobots (Fig. 1A). The principle underpinning the spatial selectivity is that a superposition of an RMF and a magnetostatic field leads to the suppression of the rotational character of the net field (**Fig. 1B**). The two representative examples shown in Fig. 1B are for the special case of an in-plane magnetostatic contribution, which shifts the circle of rotation, and for the special case of an out-of-plane magnetostatic contribution, which produces precession. At arbitrary points within a real gating field, behavior is usually intermediate between these cases.

To elucidate the transfer of torque from a superimposed RMF and gating field to microrobots, the influence of the field can first be considered numerically on the level of individual MTB. These naturally magnetic bacteria contain a stably magnetized chain of magnetite nanocrystals surrounded by a soft bacterial body that experiences viscous drag from the surrounding fluid, which we have previously modeled.^35^ **Fig. 1C** shows the angular displacement of an idealized MTB over time under increasing in-plane magnetostatic contributions in a 20 mT RMF, with a similar simulation for a 12 mT RMF shown in fig. S1A. If the magnitude of the magnetostatic component is small compared to the magnitude of the RMF, the bacterium is predicted to undergo a full rotation, although the angular velocity can vary in time (Fig. 1C and S1A). As the magnitude of the gating field increases, full rotation is suppressed and the angular displacement oscillates with increasingly limited amplitude during the cycle of the RMF.

Whereas a constant magnetic torque is expected for an MTB rotating in steady state in response to an RMF in the absence of any gating field, introducing a magnetostatic component results in time dependent torque that changes sign throughout the cycle of the RMF but remains nonzero when averaged over the cycle (fig. S1B). As the magnitude of the magnetostatic component increases and full rotation is suppressed, the time averaged torque approaches zero.

Because the results obtained for models of isolated MTB do not account for their interactions with each other and with nearby boundaries, the influence of magnetostatic gating fields on time-averaged torque density delivered to MTB confined to microchannels was considered with a continuum model. In the model, torque density was delivered under various imposed magnetostatic field magnitudes and directions. Channel geometries of different scale were examined (100 μm and 1 mm), along with two distinct stepout conditions (20 mT at 24 Hz and 12 mT at 14 Hz), as shown in figs. S1C and D. When normalized to their maximum value, averaged over all imposed magnetostatic field directions, and plotted in terms of the relative magnitude of the magnetostatic field and RMF, the time-averaged normalized torque density falls onto a similar characteristic curve (**Fig. 1D**). This curve reveals the basis for spatial selection in this system, exhibiting a nonlinear dropoff of the delivered torque density as a function of the relative magnitude of the magnetostatic gating field with the RMF. Approximately 50% of the torque density is predicted to be suppressed when the magnetostatic component reaches half the magnitude of the RMF.

### Selective actuation of MTB locally targets accumulation of companion NPs

As a first step toward empirically observing the influence of combined magnetostatic gating fields and RMFs on the actuation of MTB, we performed a set of small scale proof-of-principle experiments with our microscope-compatible arbitrary magnetic field generator. Because we have previously shown that convection generated by MTB subjected to RMFs can observably increase the diffusion of companion NPs into collagen matrices^45^, we hypothesized that this effect could be spatially restricted through the superposition of a magnetostatic gating field. To test this hypothesis, a microfluidic chip was designed with five 4-mm-diameter circular chambers arranged in a cross (**Fig. 2A**). Circular contact lines^46^ were incorporated into the center of each chamber (fig. S2A), enabling the confinement of TAMRA-labeled collagen gels in the center and formation of a barrier-free interface with the surrounding liquid compartment that contained MTB and 170 nm green fluorescent NPs (Fig. 2A). As a source for the gating field, four small rectangular NdFeB magnets with opposing magnetization directions were stably held on the sides of the device by holes cut into the PDMS (**Fig. 2B**). This arrangement produced a field free region coinciding with the central chamber of the device. By varying the geometry of the permanent magnets, gating fields with different gradients (4 T/m and 8 T/m), and thus different spatial targeting resolutions were produced (Fig. 2B and fig. S2B). As an initial test of spatially restricted torque delivery, the translational velocities of MTB in the various chambers of the 4 T/m device were observed in response to an RMF at 12 mT and 14 Hz. Velocity in the targeted central chamber was compared to the surrounding off target chambers, indicating significant suppression of delivered torque density away from the field free region (**Fig. 2C**). For subsequent experiments, the device with the lower magnetostatic field gradients was selected because the resolution of its target point was sufficient for the size of the central chamber and the smaller magnets reduced interactions with the multi-axis electromagnets.

**Fig. 2.**
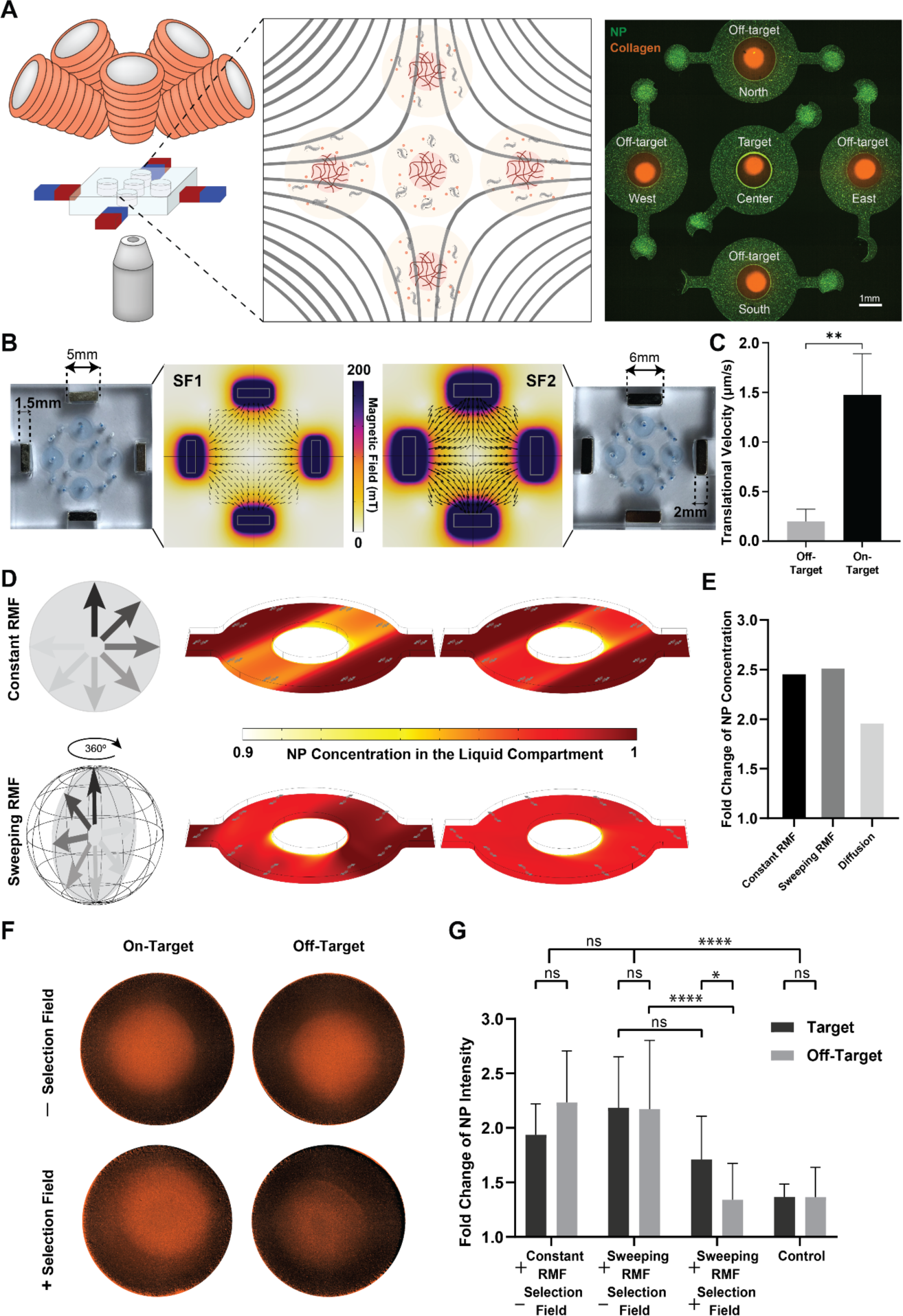
Selective actuation of MTB and localized convective effects in microfluidic devices. (**A**) Schematic of the experimental setup composed of an eight-coil electromagnetic system and a microfluidic device containing five actuation chambers, around which four permanent magnets are fixed to produce a field free point coinciding with the target chamber. The chambers offer a circular tissue-fluid interface enabled through a contact line pinning method confining collagen (red) in the core of the chambers surrounded by suspensions of NPs (green). (**B**) Computational modeling of the magnetic field generated by two differently sized NdFeB magnets. The larger set of permanent magnets generates a stronger field and thus achieves a higher spatial resolution, but also affects the working volume with higher gradient contributions (8 T/m versus 4 T/m). (**C**) Translational velocity of MTB in off-target chambers vs the target chamber, demonstrating off-target suppression (n=3 experimental replicates). (**D**) Contours of NP concentration in the liquid compartment under constant and sweeping RMF. Constant RMF leads to the emergence of a separate band inside the fluid while sweeping RMF vectors lead to a uniform increase in the transport through an effectively

Before experimentally examining the influence of focused torque delivery to MTB on companion NP diffusion, concentration profiles within the liquid compartments were modeled numerically. The results, shown in **Fig. 2D** and fig. S3, indicate that one important consideration in this experiment is the choice to either keep the plane of rotation of the RMF fixed in a single out-of-plane direction relative to the device or to sweep the plane of rotation continuously through the full circle of all possible out-of-plane rotations. A fixed plane of rotation is predicted to lead to nonuniform concentration profiles in the liquid compartment, since sustained convection in a particular direction breaks the symmetry of the device (Fig. 2D). This effect should be mitigated by continuously varying the direction of rotation, and more uniform distribution of the MTB throughout the device is also expected to modestly increase overall NP accumulation in the collagen.

Experimentally, both a sweeping RMF and fixed RMF were employed so that their influence on NP diffusion into the central collagen matrix could be compared. For the diffusion experiments, in contrast to Fig 2A, unlabeled collagen was used in the center of the chambers and 200 nm red fluorescent companion NPs placed in the surrounding compartment, so that fluorescence intensity could be used to infer concentration before and after the actuation with minimal crosstalk (**Fig. 2F**). Quantification of NPs present inside the collagen area showed a significant increase in transport under magnetic actuation for both sweeping and constant RMF in the absence of a selection field compared to a negative control relying only on passive diffusion (**Fig. 2G**). When a selection field was applied, NP transport into the off-target chambers was reduced to a level comparable to passive diffusion, whereas more than 40% of the magnetically enhanced transport was maintained within the target region. One possible source of the reduction in transport at the target point is the role of gradient forces within the experimental setup. Whereas the magnetostatic field applied here is intended primarily to extinguish torque density, its associated gradients can also apply forces to MTB that draw them to the boundary of the chambers and interfere with rotational actuation. Since the gradients in this miniaturized proof-of-concept setup are approximately twice that of the highest gradient realized in the larger setup described in a later section, these forces are expected to play a more dominant role here.

### Focused torque delivery selectively increases tumor spheroid colonization of MTB

As a physiologically relevant model of cancer, the miniaturized setup described in the previous section was adapted to study whether a magnetostatic gating field could spatially control bacterial colonization in tumor spheroids (**Fig. 3A**). We have previously found that torque-based actuation of MTB increases their colonization of tumor spheroids^30^, a widely used in vitro 3D tumor model^47, 48^. Each spheroid is composed of a mass of densely packed cancer cells that mimics some of the most relevant features of actual tumors, including cell-cell interactions and oxygen and nutrient gradients. Here, human breast cancer cells, MCF-7, were used to form spheroids roughly 400 µm in diameter, which were placed in 3 mm holes punched in PDMS in a well pattern mimicking the previous device (**Fig. 3B**). Spheroids were added to the wells with 25 µL of bacterial suspension of fluorescently labeled MTB for 1 hr of magnetic actuation followed by collection, washing, and incubation for 23 hr. Bacteria penetrating a spheroid are expected to encounter additional resistance from interactions with solid components of the spheroid, and we have found this effect to be comparable to an increased effective viscous drag, both experimentally (fig. S4) and through simulations (fig. S5). Accordingly, the RMF conditions selected to actuate the MTB here were 20 mT at 14 Hz, and the selection field with the higher gradient was employed to ensure adequate resolution in selective torque delivery.

**Fig. 3.**
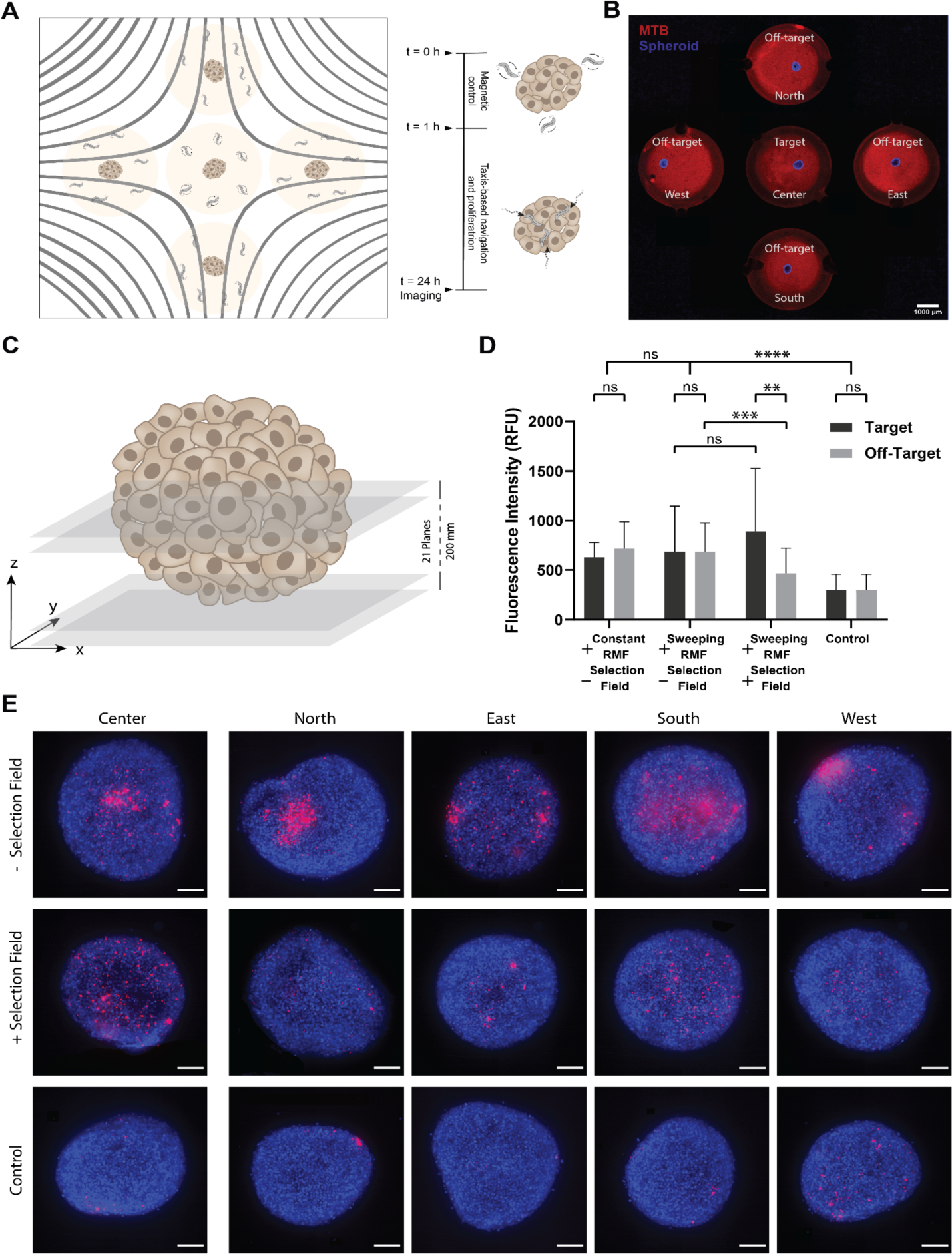
Focused torque delivery enhances tumor colonization of MTB in target locations *in vitro*. (**A**) Schematic of the experimental setup using the same configuration as above, but five open wells were provided instead of closed chambers to allow for convenient placement of 3D tumor spheroids cultured from MCF-7 cells. (**B**) Spheroids (blue) surrounded by a dispersion of MTB (red) labeled with a NIR proliferation dye allowing to track daughter cells upon proliferation. (**C**) Schematic of the readout procedure integrating the fluorescent intensity of Z-stacks at intervals of 10 µm for a distance of 200 µm along the z-axis. (**D**) Results of the analysis showing suppressed colonization in off-target spheroids compared to target spheroids (n=3, biological replicates for all conditions). Sweeping RMF consistently yielded a higher colonization than constant RMF. (**E**) Representative fluorescent micrographs of projected z-stacks for each experimental condition.

After actuation and incubation, confocal fluorescence microscopy was used to analyze the accumulation of MTB within the spheroid, with Z-stacks taken at 10 µm intervals up to a total height of 200 µm (**Fig. 3C**). The MTB had been labeled with a far red proliferative dye to ensure that daughter cells resulting from proliferation during incubation could also be detected. In the absence of a selection field, significantly higher accumulation (2.3 fold) was observed after exposure to sweeping RMFs compared to the unactuated control (**Fig. 3D**). With the introduction of a selection field, MTB colonization of the on-target spheroids did not drop significantly, whereas a significantly suppressed accumulation was observed at off-target sites in the presence of the superimposed gating field (Fig. 3D). Clusters of bacteria inside the tumor spheroids were visible after 24 hours of incubation, as shown in the representative Z-projected confocal images of the tumor spheroids depicted in **Fig. 3E**.

### Design and validation of a mouse-scale setup for focused torque application

The in vitro proof-of-concept experiments described in the previous two sections demonstrated the feasibility of combining selection fields with RMFs for spatially restricted torque-based actuation. To take steps toward testing these concepts in vivo and exploring potential form factors for scaled up instrumentation, a mouse-scale setup was designed and constructed. Firstly, the working volume of the RMF should encompass the entire animal and its magnitude should be in the range of 10 to 20 mT for comparability to the in vitro experiments. Next, the volume of the desired field free region formed by the selection field should be approximately 1 cm^3^ in volume to coincide appropriately with a flank tumor. Lastly, the ability to manipulate the position of the field free region with respect to the mouse was also desired to ensure that it could be accurately targeted to the tumor.

An exploded view of the design that was developed to meet these requirements is shown in **Fig. 4A**, along with its assembled form including a representation of an anesthetized mouse in **Fig. 4B**. It consists of three main components: 1) an array of permanent magnets that produces a field free point (fig. S6), 2) a set of AC coils that are used to generate the RMF (fig. S7), and 3) a DC offset coil that moves the zero point within the transverse plane of the animal and rejects heat through forced water cooling (fig. S8).

**Figure 4.**
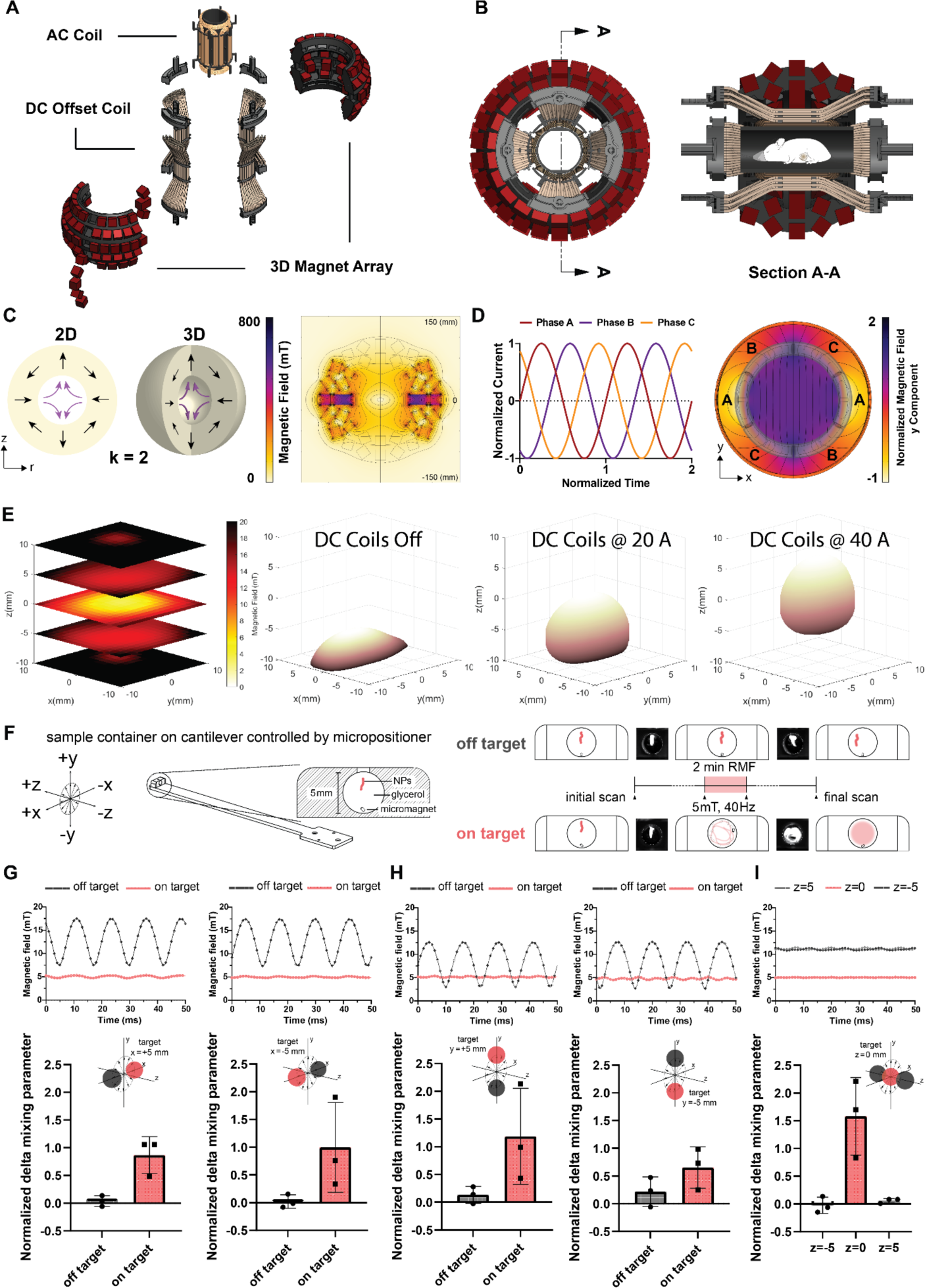
Characterization and validation of a mouse-scale torque focusing apparatus. (**A**) Exploded 3D view of the device, showing the three main components labeled. (**B**) Assembled 3D view of the device, shown in cross section with a representation of a mouse. Inner diameter of the working volume is 5.5 cm. (**C**) The concept of a 3D “magic sphere” geometry is derived from an axisymmetric revolution of a second order Halbach cylinder (left). A finite element model indicates the expected magnitude of the field produced by the modified geometry actually used (right), which excluded magnets from the poles to preserve access and used a combination of ferrite ceramic and neodymium iron boron magnets to improve isotropy of the zero point. (**D**) 120° phase shifted input currents are used to drive the three-phase RMF coil and generate a sufficiently uniform rotating magnetic field. A finite element model of the field generated by three phase stators around a transverse section of the RMF coil shows the expected uniformity of the field. (**E**) An array of 64 Hall probes was used to measure and reconstruct the field free region of the permanent magnet array. Currents in the DC offset coils displace the position of the field free region, with measured field data represented here in terms of the interpolated surface of points at 7.5 mT field magnitude. (**F**) A schematic of the experimental design of the micromagnetic mixing assay used to validate selective torque-based actuation in the full setup. Representative examples of initial and final scans showing the distribution of fluorescent NPs are inset. (**G**)-(**I**) Hall probe measurements showing field magnitude (above) and normalized changes in mixing parameter (below) are provided for target points at x = ±5 mm in (**G**), y = ±5 mm in (**H**), and z= 0, ±5 mm in (**I**). “On target” for the x and y axis refers to offset currents that bring the target point into alignment with the mixing chamber at the respective points. “On target” for the z axis refers exclusively to position at zero offset current.

The array of permanent magnets responsible for generating a magnetostatic gating field was based on a “magic sphere” geometry, formed by an axisymmetric revolution of a second order Halbach cylinder^49^ (**Fig. 4C**). The desired distribution of magnetization was approximated with 2 cm X 2 cm X 2 cm stacks consisting predominantly of ferrite ceramic permanent magnets (grade Y35) affixed to a 3D printed frame that held them at the appropriate angles. To ensure access to the working volume, magnets were omitted from the poles of the magic sphere, and sintered neobymium iron boron magnets (N45) were strategically incorporated around the equator of the array to improve the isotrophy of the filed free point. The results of an axisymmetric finite element model show the expected distribution of magnetostatic field magnitude, with a central field free region that is an oblate spheroid with its shortest axis coinciding with the axis of symmetry.

Strong RMFs are routinely generated at the scale of tens of centimeters for industrial applications such as 3-phase industrial motors, which acted as a source of inspiration for the RMF coil geometry in this setup. Three independent stators were wound around a 3D printed frame and driven with 3-phase current to produce an adequately homogenous RMF, with its plane of rotation perpendicular to the axis of the cylindrical working volume (**Fig 4D**). The target field magnitude of 20 mT was achieved in the 5.5 cm diameter, 8.5 cm long working volume, but continuous operation was limited in practice to 15 mT based on heat rejection capabilities.

The two DC offset coils were arranged in the same basic concentric geometry as the RMF coils, residing between the RMF coils and the permanent magnet array (Fig. 4A). These coils were formed manually from 3 mm copper tubing and arranged geometrically using a set of 3D printed guides and flow splitting manifolds (fig. S8) Electrical current was run in series through these coils, whereas chilled water flowed in parallel to cool the setup to temperatures safe for anesthetized rodents. Thermal contact between the RMF coils and the DC offset coils was made via electrically non conductive thermal epoxy.

The obtained field free point was spatially characterized with a fixed array of 64 individual Hall probes, with its ellipsoidal shape found to be in good agreement with expectation (**Fig. 4E**, fig. S9). This array was then also used to observe the displacement range of the field free point by introducing currents into the DC offset coils, depicted in **Fig. 4D** by the electronically controlled motion of an interpolated surface of constant magnitude at 7.5 mT. As expected from simulation (fig. S10), approximately 1 cm of travel in the transverse plane was possible for sustained offset with this setup, ultimately limited in practice by heat dissipation of the uncooled portions of the DC coils.

Final characterization of the net magnetic fields produced by the fully assembled setup (fig. S11) was conducted in tandem with a NP mixing experiment intended to validate spatially selective torque-based actuation. In brief, a cantilever was affixed to a micropositioner, which was used to maneuver a Hall probe at the center of the field free point, enabling measurements at known positions, combined with the RMF and DC offsets. Keeping the same coordinate reference, a 3D printed 5 mm diameter cylindrical sample chamber sealed by clear polycarbonate sheet and filled with glycerol was placed at the end of the cantilever and introduced back into the setup at predetermined locations (see **Fig. 4F**). This chamber also contained a microscale cylindrical neodymium magnet (300 μm diameter X 500 μm length) freely situated at the bottom, as well as a localized 1 μL injection of red fluorescent NPs toward the top. Unsuppressed rotational actuation of the micromagnet at conditions near its stepout frequency (40 Hz at 5 mT in pure glycerol), caused it to move rapidly throughout the well and convectively mix the NPs with the surrounding glycerol. RMF suppressed by a magnetostatic contribution resulted in oscillation of the micromagnet at the bottom of the well, with minimal contributions to mixing. The degree of mixing before and after actuation was assessed by examining changes to the mixing index (a function of the distribution of fluorescence intensity within the chamber) before and after actuation, with representative examples of the scans provided in Fig. 4F.

To test the spatial selectivity of the zero point in each direction, Hall probe measurements and mixing experiments were performed at paired displacements in the transverse plane for x and y = ±5 mm from the zero point, with the DC coil currents adjusted to bring the field free point into alignment for “on target” cases (**Fig. 4G, 4H**). These results indicate spatially selective torque-based actuation in the mouse scale setup, and moreover confirm the functionality of the DC offset coils. Spatial selection along the axis of the cylinder (i.e. in the z direction, **Fig. 4I**) is accomplished through positioning rather than through electronic control of DC currents. Thus, the Hall probe characterization was repeated along this axis at three points, z = −5 mm, 0 mm, and +5 mm, in order to verify similar functionality in this direction (Fig. 4I).

## DISCUSSION

It has previously been asserted that magnetic torques rather than magnetic forces are advantageous for actuating the smallest microrobots to efficiently deliver mechanical energy via externally applied magnetic fields. The present work illustrates a further advantage of torque-based schemes–their compatibility with an open loop control strategy for spatially restricted transmission of torque density, based on combining RMFs with magnetostatic gating fields. One way to view the mouse scale setup that was developed and validated here is as an unusual type of three-phase motor with numerous diffuse and distributed rotors, accompanied by a mechanism to exert spatial control over which of these rotors are effectively locked and which are free to rotate. One clear alternative approach to achieve selective rotational actuation is to apply spatially limited RMFs, a strategy that was indeed exploited in our previous work^30^. Nevertheless, the introduction of a gating field more tightly and controllably restricts a focal point for torque delivery, in addition to permitting target points that are remote from the coils or electromagnets generating the RMF.

Although the use of gating fields in this work was inspired by the mechanism for signal isolation employed in MPI, key differences should be noted. Firstly, the resolution of field free regions in MPI depends directly on material properties of the magnetic tracer NPs, whereas the resolution here is set primarily by the magnitude of the RMF relative to the gating field gradient. Microrobot design thus influences resolution of the delivered torque density only indirectly, because magnetic and hydrodynamic characteristics do ultimately dictate optimal RMF conditions. Secondly, in MPI, gating fields with higher gradients are almost always desirable to improve resolution, aside from the technical difficulties associated with generating them. In the case of focused torque delivery, gradients need to remain low enough to avoid exerting forces that pull microrobots against surfaces and interfere with their rotation, an effect that may have played a role in the miniaturized in vitro experiments presented here. This implies that for each microrobot design, there exists both a minimum gating field gradient needed to provide the required resolution, as well as a maximum gradient that can be tolerated before interfering with torque-based actuation. Fortunately, for the subset of microrobots small enough to be suitable for intravenous administration (typically less than 10 μm) like the MTB employed here, and for gradients that can be realistically generated at physiological scales, the influence of torque typically far exceeds that of gradient forces.

While it would likely be possible to preserve the basic geometry of the in vivo setup developed here while scaling it up to the size of human patients, more efficient approaches are readily foreseeable. Our in vivo experiment specifically required an apparatus capable of exposing the whole body of a mouse to an RMF, whereas this requirement could be relaxed for a clinical instrument dedicated to focused torque delivery. Constructed around the working distance as a key scaling parameter, a comparatively smaller permanent magnet array could be employed, possibly with mechanical adjustability for zero point focusing. Our early in vitro experiments showed functional benefits to using swept RMFs, and an ideal clinical apparatus should also be capable of sweeping its plane of rotation. Finally, compatibility with imaging modalities such as ultrasound or computed tomography in order to initially pinpoint target coordinates is likely an important consideration informing the design of a clinically relevant device.

The concept of focused delivery of torque density to biomedical microrobots could eventually be applied to numerous magnetic microrobot designs. The underlying logic of torque suppression using gating fields may even prove applicable to non-rotating torque-based locomotion schemes, such as those employing beating magnetic fields, a topic that could warrant further investigation. Many different apparatuses can also be envisioned for focused torque-based actuation, ranging from altered use cases of existing MPI systems to specialized devices that incorporate permanent magnet arrays. As an open loop control strategy, torque focusing is appealing because it brings together the simplicity and passivity of a defined target point with a scheme that powers the activity of biomedical microrobots.

## MATERIALS AND METHODS

### MTB culture and fluorescent labeling

Cultures of *Magnetosprillum sp.* AMB-1 (ATCC® 700264™) were used for this study. Bacteria were routinely passaged after 7 to 10 days and grown using magnetic spirillum growth medium (MSGM, ATCC® Medium 1653). Cultures were kept at under anaerobic conditions in 15 ml tubes.

To stain the bacteria and allow tracking of subsequent generations, 2 µl of a farred proliferative dye (CellTrace™ CFSE Cell Proliferation Kit, ThermoFischer) was added to 1 ml of bacteria suspension at 5×10^8^ cells/ml, assessed by optical density measurements. Following 20 min of agitation on a shaker while protected from light, the dye was deactivated using 100 µl of DMEM for 10 min. The bacteria were spun down and resuspended in 1 ml DMEM for subsequent experiments.

### Selective actuation of MTB with NP transport experiments

An experimental platform featuring five separate microfluidic wells (North, East, South, West, Target) made from polydimethysiloxane (PDMS) was fabricated according to standard soft lithography protocols. Each well features a circular ridge to allow surface tensions based contact line pinning of viscous collagen (**Supplementary Figure S2**). After 2-3 days after plasma bonding, the microfluidic devices were fitted to hold four permanent magnets (top, bottom, left, right) within precut slots for accurate positioning and creation of a FFR. To generate a desired gating field with FFR in the center well (Target), a simplified version of the Halbach cylinder k=2^49^ was adopted using NdFeB block magnets (5 x 2.5 x 1.5 mm or 5 x 2.5 x 1.5 mm, N45, Supermagnete.ch). At the top and the bottom, arbitrarily defined north poles faced each other while north poles pointed outward on the left and the right sides. Upon assembly, collagen was filled into the central part of each well. Following 45 min incubation at 37 °C, the devices were stored in humidified petri-dishes at room temperature before the start of the experiment. MTB were spun down and resuspended in PBS at a final concentration of 8.7×10^9^ cells/ml. To quantify NP transport, 1 µl of red fluorescent NPs (FluoSpheres carboxylated microspheres, 0.2 µm) were added per 100 µl of the MTB suspension. The final suspension was then introduced into the annular part of the wells surrounding the central collagen compartment, followed by torque-based magnetic actuation for 1 hour. Transport evolution was monitored and captured by confocal micrographs at t= 0, t= 30 min and t=1 hour.

Image processing was performed in ImageJ (NIH). The collagen area was determined via the Analyze Particles command applied to the binarized image of the red channel. Fluorescent signal from NPs inside this ROI was integrated for different time points. The values of later time points were normalized by the time point 0 to compensate for any initial penetration of the NPs caused by the filling process. The ROI defined by the collagen was shrunk to form bands at certain intervals which enabled quantification of the signal as a function of the distance from the interface.

### Magnetic actuation in small proof-of-concept setup

The RMF was produced in these experiments (shown in Fig. 2 and 3) with a commercial electromagnet setup consisting of 8 coils forming a hemisphere at one end as the working space was used for all magnetic actuations (MFG-100-i, Magnebotix AG). The setup was integrated into an inverted microscope (Nikon Ti Eclipse), enabling live imaging of the samples under magnetic actuation. The system can generate uniform arbitrary 3D magnetic fields within 1 cm^3^ in space. Sweeping RMF was applied through application of the following input functions:

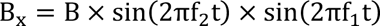

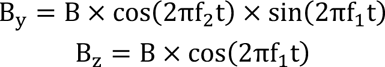

Where f_1_ is the frequency of the out-of-plane RMF and f_2_ represents the sweeping frequency of the plane of rotation. In order for the plane of rotation to go through one revolution during the actuation, f_2_ was set to 1/3600. Under the especial case of f_2_=0, the actuation scheme turns into constant RMF with yz as the plane of rotation. For experiments with the alternating RMF, one plane of rotation, i.e. yz, with switching axis of rotation between x and -x directions were used.

### Spheroid culture

MCF-7 cells were cultured in high glucose Dulbecco’s Modified Eagle’s Medium (DMEM, ThermoFischer) supplemented with 10% FBS and 1% P/S. Ultralow adhesion plates (InSphero) were utilized to form tumor spheroids from MCF-7 cells. A seeding density of 10,000 cells/well in 50 µL of growth media was used in 96 well plates. The well plates were centrifuged at 500 x g for 10 min followed by incubation at 37 °C with 5% CO_2_. The size of the tumor spheroids reached ∼400 µm after 3 days. DNA staining with Hoechst 33342 at a final concentration of 5 µg/mL in media was performed before the experiment for one hour at 37°C.

### *In vitro* experiments under torque focusing

For these experiments, the five-chamber devices were punched at the center of each well with 3 mm punchers resulting in five wells for placing the spheroids. Corresponding slots for small magnets were cut and magnets were placed following the above-mentioned order for magnetization directions. Wells were rinsed with ethanol, air dried, and then exposed to UV for an hour for sterilization. Each well was filled with ∼25 µl of MTB suspension, followed by careful release of spheroids and settling at the bottom of the wells. The devices were then covered with a lid and mounted on the microscope holder to minimize contact with the surrounding air and avoid potential contaminations. Magnetic actuation at 20 mT and 14 Hz was applied for 1 hour,followed by collection of the spheroids, through washing with media, and incubation in sterile well plates at 37°C and 5% CO_2_ for 24 hours.

Spheroids were transferred into PDMS pools, bonded on thin cover slips for high resolution confocal imaging after 24 hours. Images were taken with along the Z-axis for a distance of 200 µm at 10 µm intervals. To quantify the accumulation of bacteria in tumor spheroids, the ROI was defined as the outer edge of the spheroid in a binarized Z-projected image of the DAPI channel. The far-red signal was then integrated for all planes according to the respective ROI.

### Computational modeling of magnetic actuation and NP transport

The “Magnetic Fields No Currents” physics interface of COMSOL Multiphysics was utilized to model the magnetic field distribution inside the working space of the microfluidic devices and resulting transport of NPs therein. Four blocks representing the respective permanent magnets were created according to their experimental layout the size of the blocks and their mutual distances were varied in a parametric study. Each magnet was modelled as a permanent magnet possessing a remnant flux density of 1.4 T and a recoil permeability of 1.05.

To study NP transport, the geometry of a single well from the multi-chamber design was recreated. Governing equations from Stokes Flow and Transport of Diluted Species interfaces were solved for fluid velocity and concentration of the particles. The effect of magnetic actuation on MTB was modelled through application of a volume force corresponding to the torque-driven flow of bacteria in such chambers. The diffusion coefficient of the NP in liquid was calculated from the Stokes-Einstein relation (D_0_ = 2 × 10^−12^ m^2^/s). Collagen gel was modelled as a porous material with porosity of 0.6. Hydraulic permeability was assumed to be *k* = 10^−10^ m^2^ and diffusion coefficient was calculated by modifying the fluid diffusion coefficient according to the tortuosity model for effective diffusivity.

## Supplementary Material

**Figure S1.**
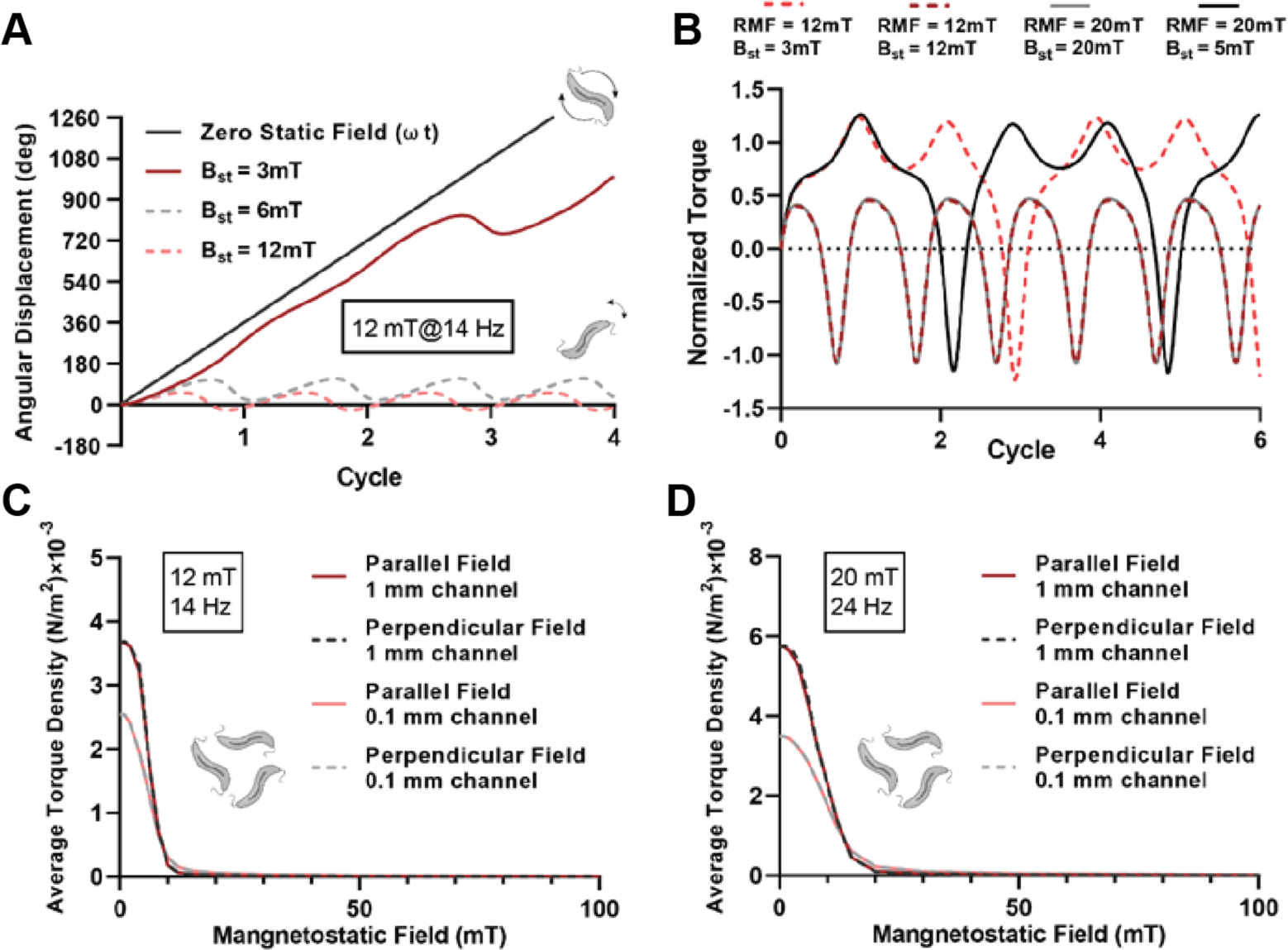
Selection field requirements for suppression of torque-driven transport. (**A**) Angular displacement of individual microrobots exposed to different magnitudes of the static gating field with RMF 12 mT and 14 Hz. (**B**) Applied magnetic torque on a single microrobot under various RMF and gating field values. Average torque drops to almost zero reflecting the transition from rotation with back-and-forth motion to symmetric oscillation. (**C, D**) Decay of the average torque density across channels of varying widths as a function of differently oriented magnetostatic fields under RMF of (**C**) 12 mT and 14 Hz and (**D**) 20 mT and 24 Hz.

**Figure S2:**
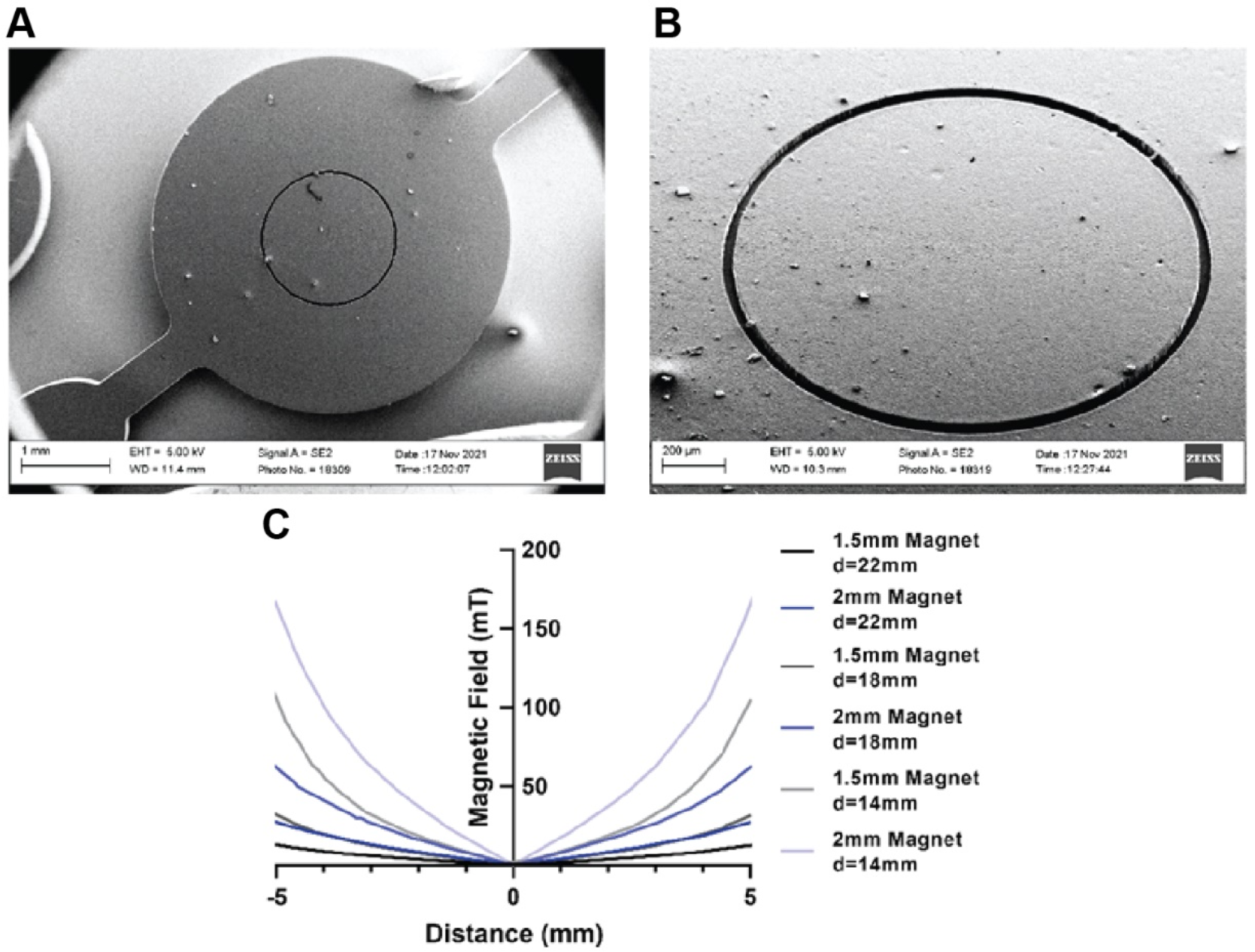
Design and characterization of the in vitro platform to study selective actuation of MTB and selective transport of NPs. (**A**) SEM micrograph of the microfluidic master mold from top for the device shown in Fig 2A of the main text. (**B**) SEM micrograph of the microfabricated ridge in a ring shape serving as the contact line to pin viscous collagen upon filling into the chambers. (**C**) Magnetic field profiles throughout the working volume of the device layout shown in Fig 2B of the main text, as predicted by finite element modelling assuming two differently sized small blocks of NdFeB magnets at different distances. Small magnets at d=18 mm and large magnets at d=22 mm provide adequate resolution for selective actuation of target well.

**Figure S3:**
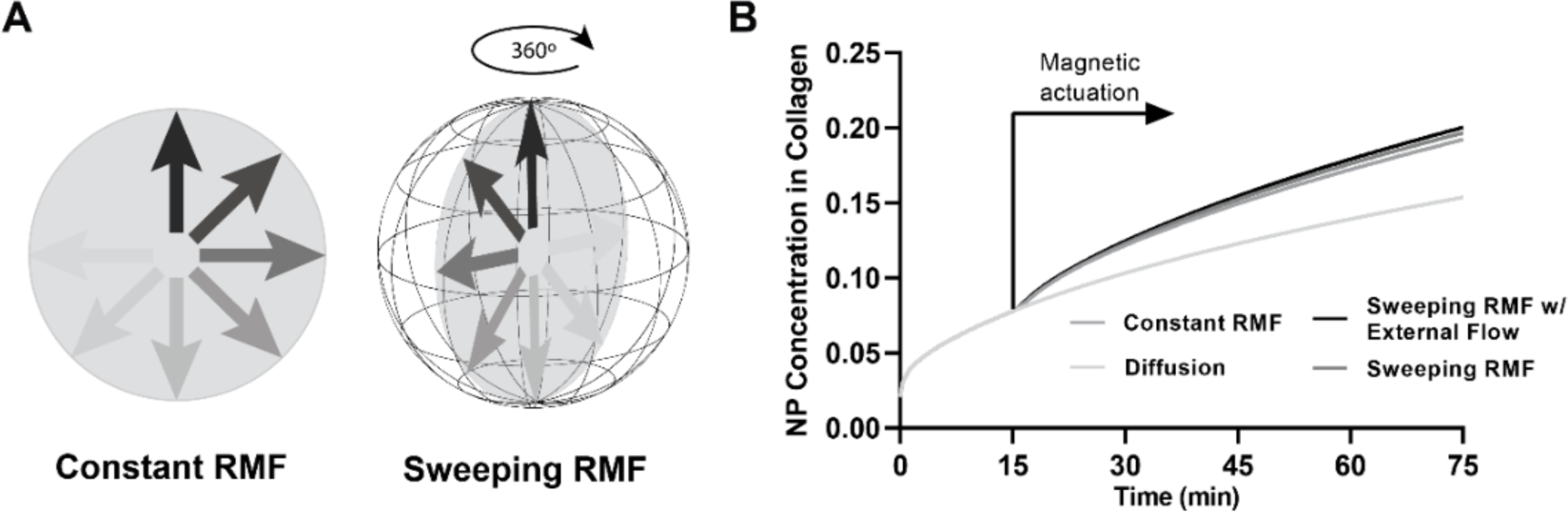
Simulated magnetically enhanced selective transport of NPs into collagen gels. (**A**) Schematic of sweeping RMF as an actuation scheme for isotropic targets. Plane of rotation undergoes one revolution during the experiment with sweeping RMF opposed to constant RMF where the plane of rotation is fixed during the actuation period. (**B**) Computationally resolved time evolution of NP concentration in collagen modelled as a porous material. Sweeping RMF slightly outperforms constant RMF, and presence of fluid flow inside the chambers has subtle effect on increase of the transport.

**Figure S4.**
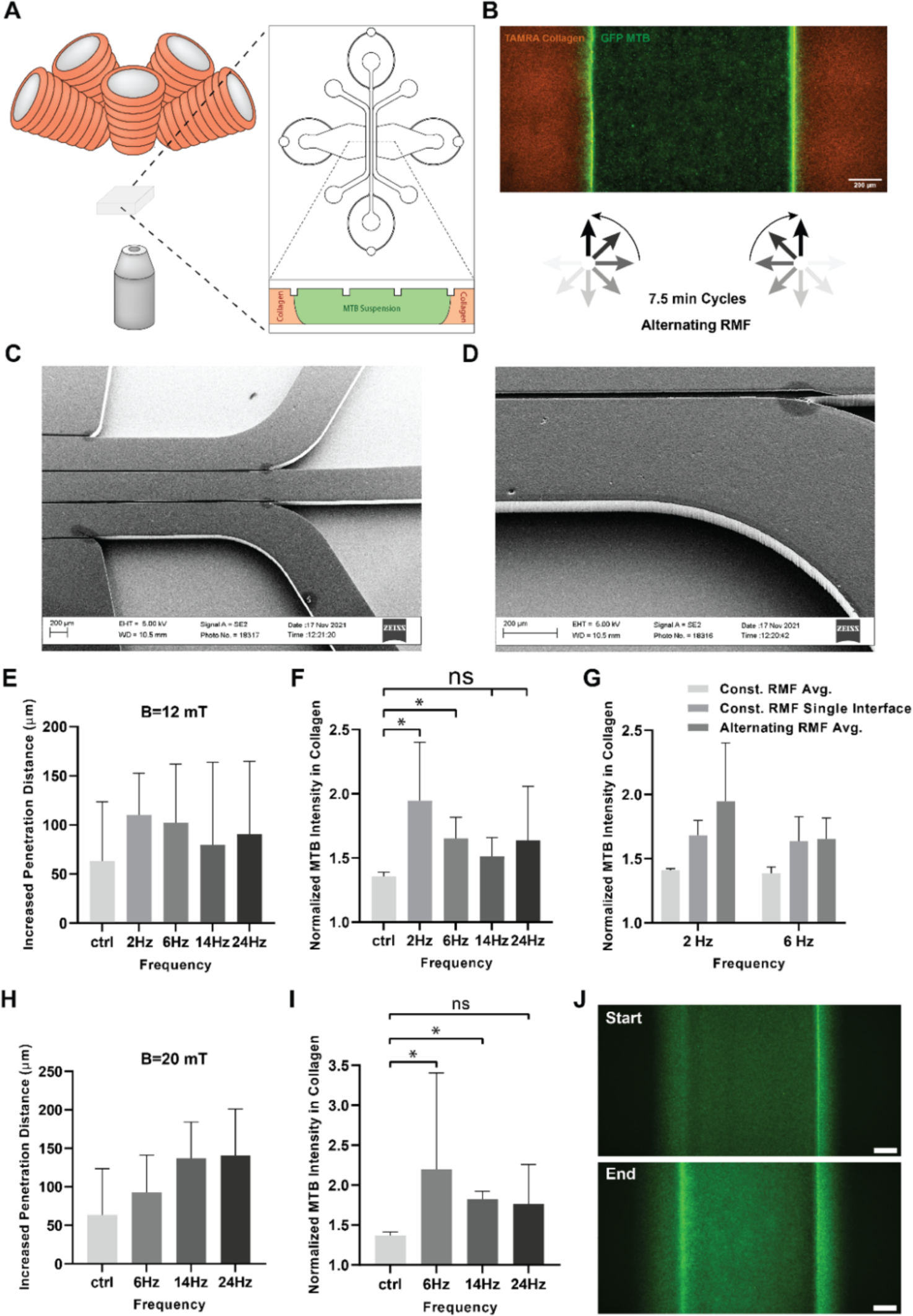
In vitro test platform to study influence of RMF driving condition on MTB penetration into tissue mimicking matrices. (**A**) Schematic of the contact line pinning microfluidic devices exposed to magnetic actuation. (**B**) Compartmentalization of the bacteria and collagen gel through microfabricated contact lines. Suspension of GFP-expressing bacteria (green) is introduced between two chambers filled with TAMRA labelled collagen (red). Alternating RMF was used for balanced penetration of the bacteria at both interfaces. (**C,D**) SEM micrographs of the microfabricated ridge in a ring shape acting as the contact line. (**E**) Increased penetration distance over the course of the experiments (30 min) under RMF at 12 mT with different frequencies. (**F**) Integrated MTB signal in collagen at the end of the magnetic actuation sequence at 12 mT, normalized by the initial timepoint. (**G**) Comparison of MTB penetration under different actuation schemes. Targeted single interface at constant RMF exhibits similar values to the average of both interfaces under the alternating RMF. (**H**) Increased penetration distance over the course of the experiments (30 min) under RMF at 12 mT with different frequencies. (**I**) MTB signal in collagen at the end of the magnetic actuation at 12 mT normalized by the initial time point. (**J**) Representative images of the bacteria before and after the magnetic actuation (scale bars: 200 µm)

**Figure S5.**
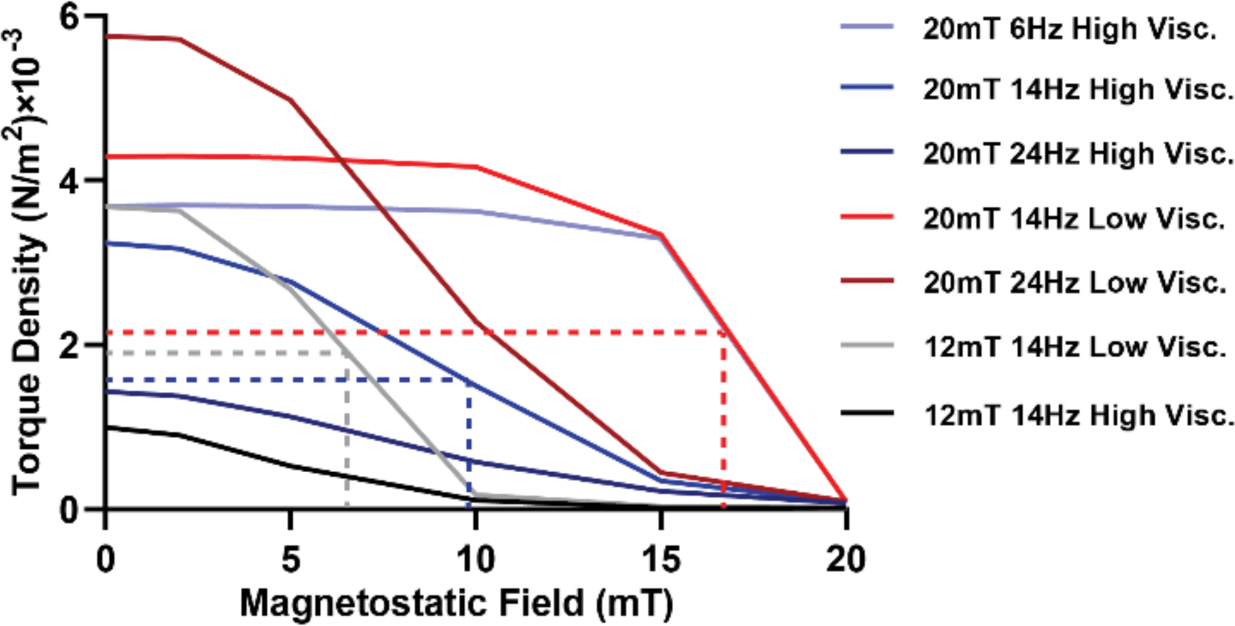
Simulated average torque densities in bacterial suspensions under various RMF parameters in low and high viscosity mediums. Increasing the field magnitude in the presence of higher resistance compensates for the loss in torque density under weaker fields and increases the required magnetostatic field to suppress off-target transport.

**Figure S6.**
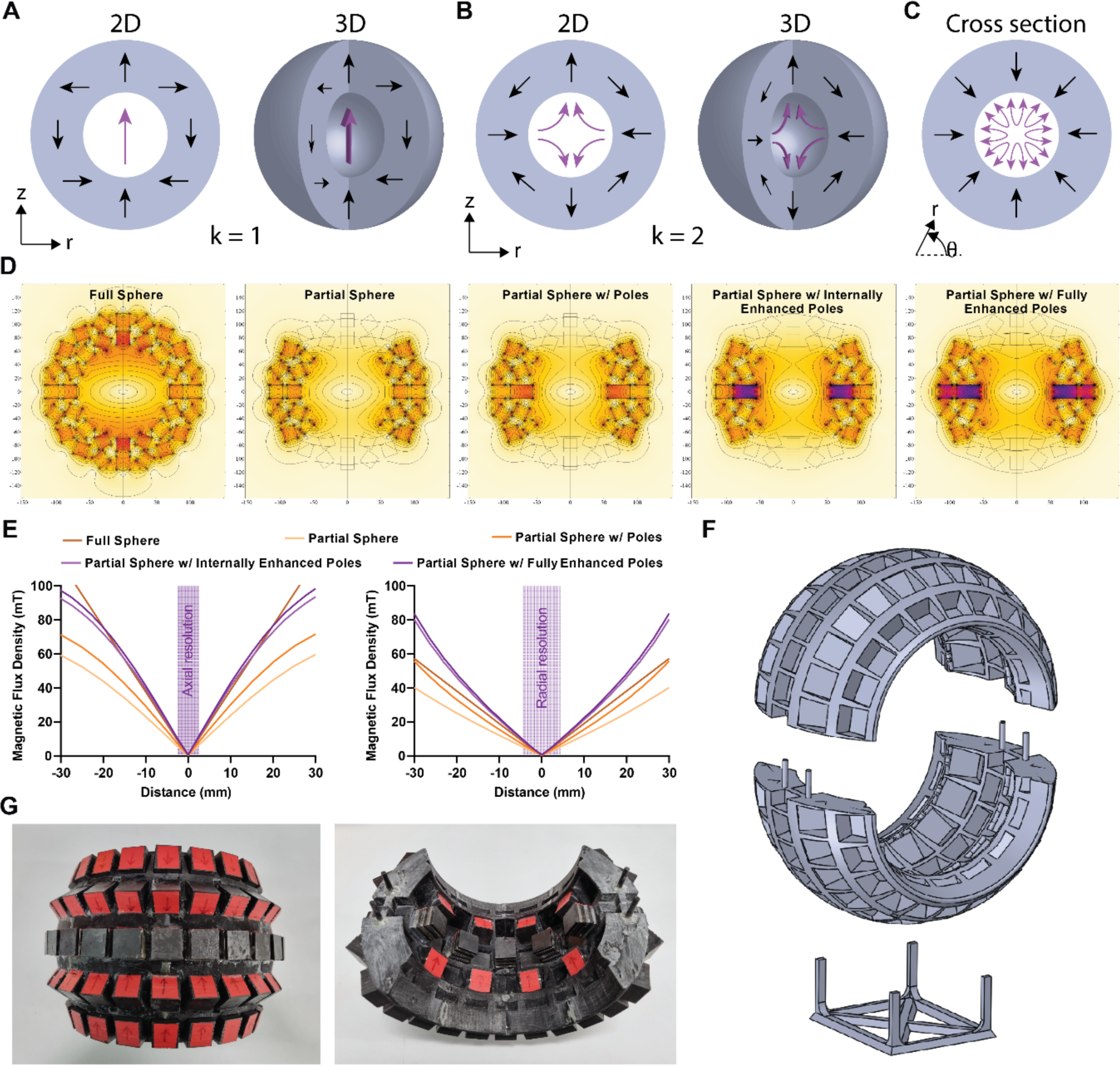
Description and simulation of magic sphere array to generate the desired magnetostatic selection field. (**A**) Magic sphere representing the transformation of Halbach cylinder k=1 into 3D for uniform magnetic fields. (**B**) 3D transformation of the Halbach cylinder with k=2 to create a zero point in 3D space. (**C**) Cross section of the sphere resulting from revolution of the k=2 array. (**D**) Computational modeling of the magnetic field generated by the k=2 magic sphere formed by ferrite block magnets. Geometric constraints from the DC coil lead to a partial sphere with somewhat reduced resolution. Adding additional magnets to the middle row and supplementing them with stronger NdFeB magnets restores the lost confinement of the zero point. (**E**) Simulated magnetic field profiles in axial and radial directions. Higher resolution in the axial direction arises from deviation of the array in cross sectional plane from ideal k=2 arrangement. Profiles confirm the desired 1 cm resolution was attained by creating enhanced poles in the middle row of the sphere. (**F**) Designed geometry for the skeleton of the modified magic sphere. Indentations on both inner and outer surfaces are incorporated as slots for stacks of block magnets. (**G**) Fabricated modified magic sphere with mounted magnets. Inner and outer layers of magnets are assembled in way that ensures 2-fold symmetry of the zero point, assuming both halves are brought together fully. In practice, the gradient was somewhat stronger in the y direction than the x direction due to a gap that remained after clamping both halves together.

**Figure S7:**
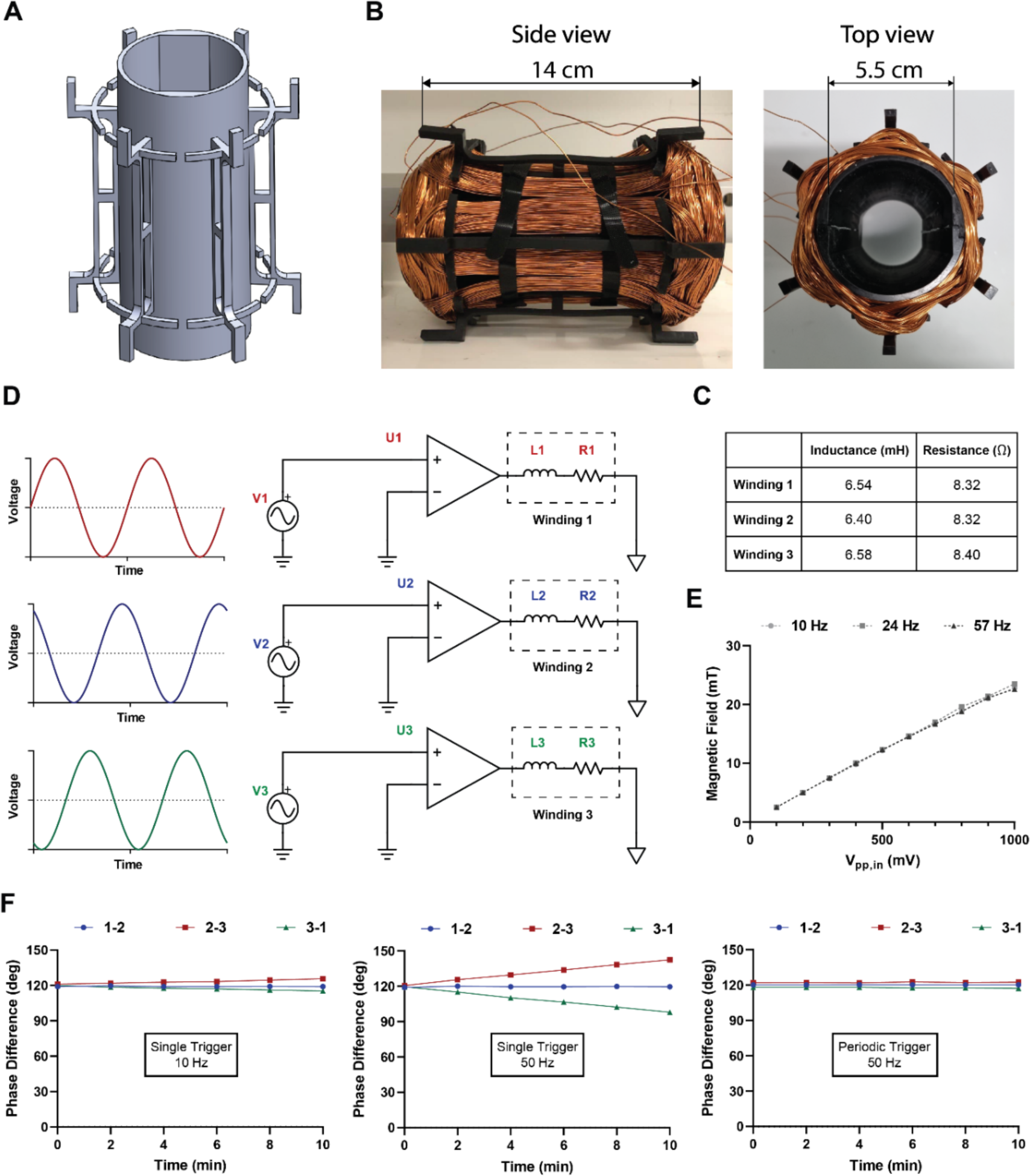
Supplementary description of the three-phase coil design for generating full-body RMF for mice. CAD geometry of the stator designed for the three-phase winding. Dedicated slots to each winding ensure compact structure of the AC coil. (**B**) Wound AC coil around the 3D printed stator. The central portion of the coil exhibits highest compactness providing space for the DC coil. (**C**) Electrical properties of individual windings forming the AC coil. Symmetric windings yield less than 3% variability in electrical properties. (**D**) Electric circuit of the RMF generating component. Phase-shifted input signals from Digilent devices are amplified and sent to the windings of the coil. (**E**) Magnitude of RMF generated by the AC coil as a function of peak-to-peak input voltage. Amplification of the signal from the waveform generators is consistent in the desired range of frequencies and fulfills the 20 mT requirement for the magnitude. (**F**) Cross-triggering strategy for synchronizing input from two waveform generators. When two waveforms are generated with a certain phase lag from two different devices, minute deviations in each cycle propagate over time and disrupt the phase different particular at higher frequencies. Periodic triggering resets the phase lags leading to consistent generation of uniform RMF.

**Figure S8:**
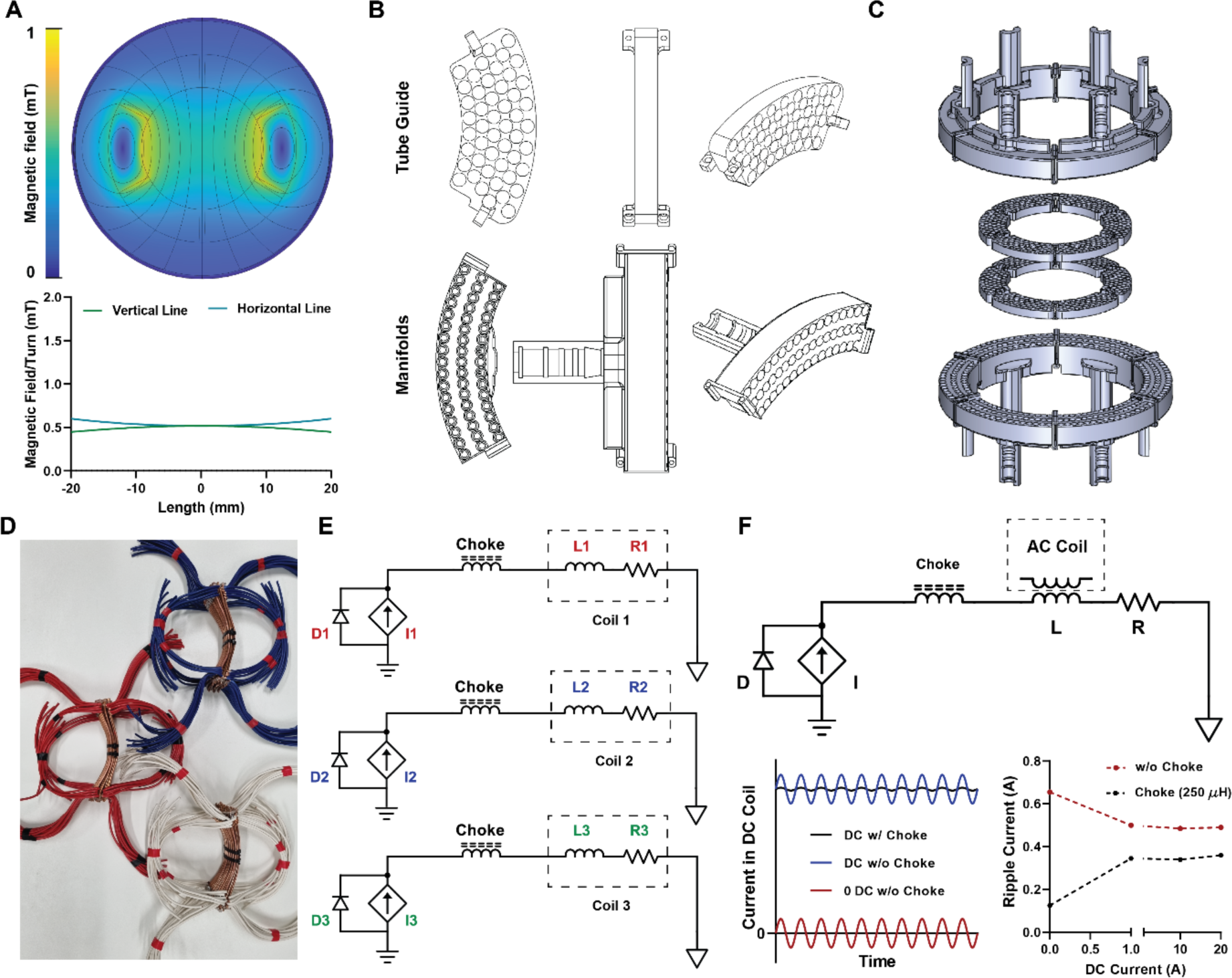
Design and fabrication of the DC offset coils. (**A**) Idealized 2D simulation of the static field generated per each turn of one coil pair. 0.5 mT per ampere-turn provides an upper bound for the attainable field from each phase. (**B**) Tube guide and manifold geometries. Tube guides serve to hold the copper tubes straight and in close contact with the AC coil. Note that an insulating laquer is applied to the copper to minimize the risk of short circuits. Manifolds are fabricated to collect the circulating water in the hydraulic circuit enabling efficient heat dissipation in the system. (**C**) CAD assembly of the tube guides and manifolds. Distances are set according to the geometrical constraints from the AC coil and the array of magnets. (**D**) Single coils composed of copper tubes and connecting wires, before their full assembly. The original DC offset coil was planned to contain 3 pairs of parallel tube asseblies, but space contraints later caused this to be revised to 2 pairs. (**E**) The originally planned electrical circuit of the system to power the DC offset coils. Chokes are included to suppress the ripple current due to inductive coupling with the RMF coils and diodes parallel to the power supplies protect them from reverse current caused by inductive reactance in the circuit. The diodes and chokes were omitted from the design actually used for the mouse scale setup because they were found to be unnecessary.(**F**) Effect of the choke in circuits with two smaller inductively coupled test coils. Reduced suppression of ripple noise at higher currents implies that large iron powder cores with lower permeability and relatively modest number of turns are required to avoid saturating the choke. These considerations would become more important if the operating frequency of the setup were raised substantially.

**Figure S9:**
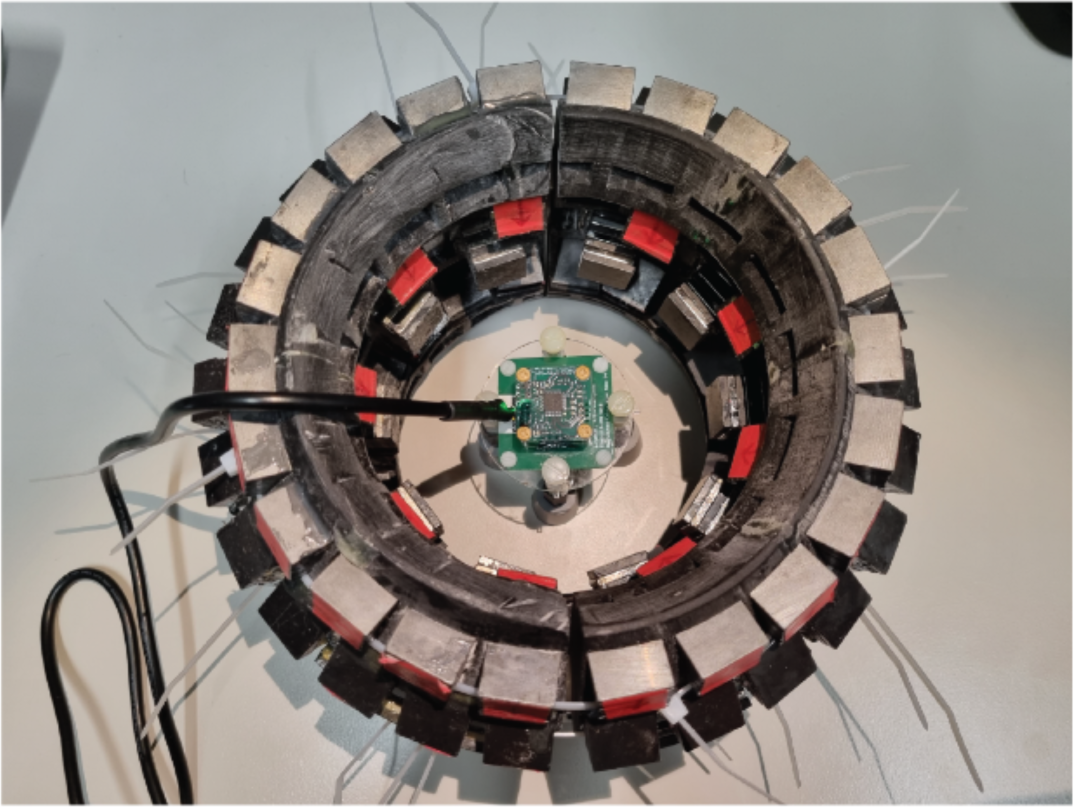
Characterization of the magnetostatic field generated by the array of permanent magnets. assembled magic sphere for is shown. A cube of Hall sensors located at the center of the sphere measures the selection field from the magnets in a 3D volume.

**Figure S10:**
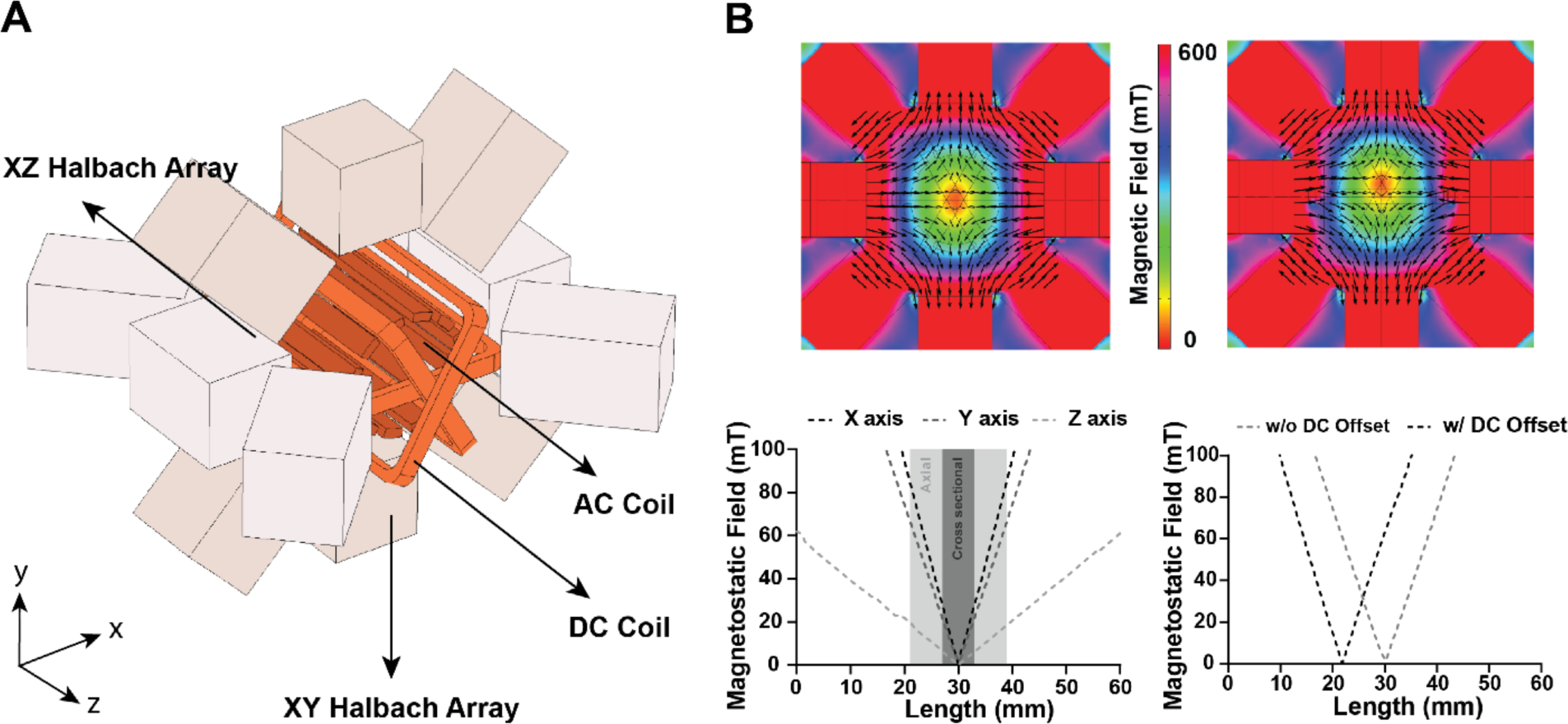
Preliminary computational study of displacement of the field free point. (**A**) Simplified multilayer configuration envisioned for the selection field setup. Note that the configuration of permanent magnets reflects an earlier approximation of the array that was ultimately used, but the concept remains the same. The DC coil positioned between the AC coil and array of magnets is responsible for adding offset to the magnetostatic field from the permanent magnets and moving the zero point. (**B**) Computational modeling of the static field from the Halbach arrays and the DC coil. Running current through the DC coils leads to shift in the zero-point location.

**Figure S11:**
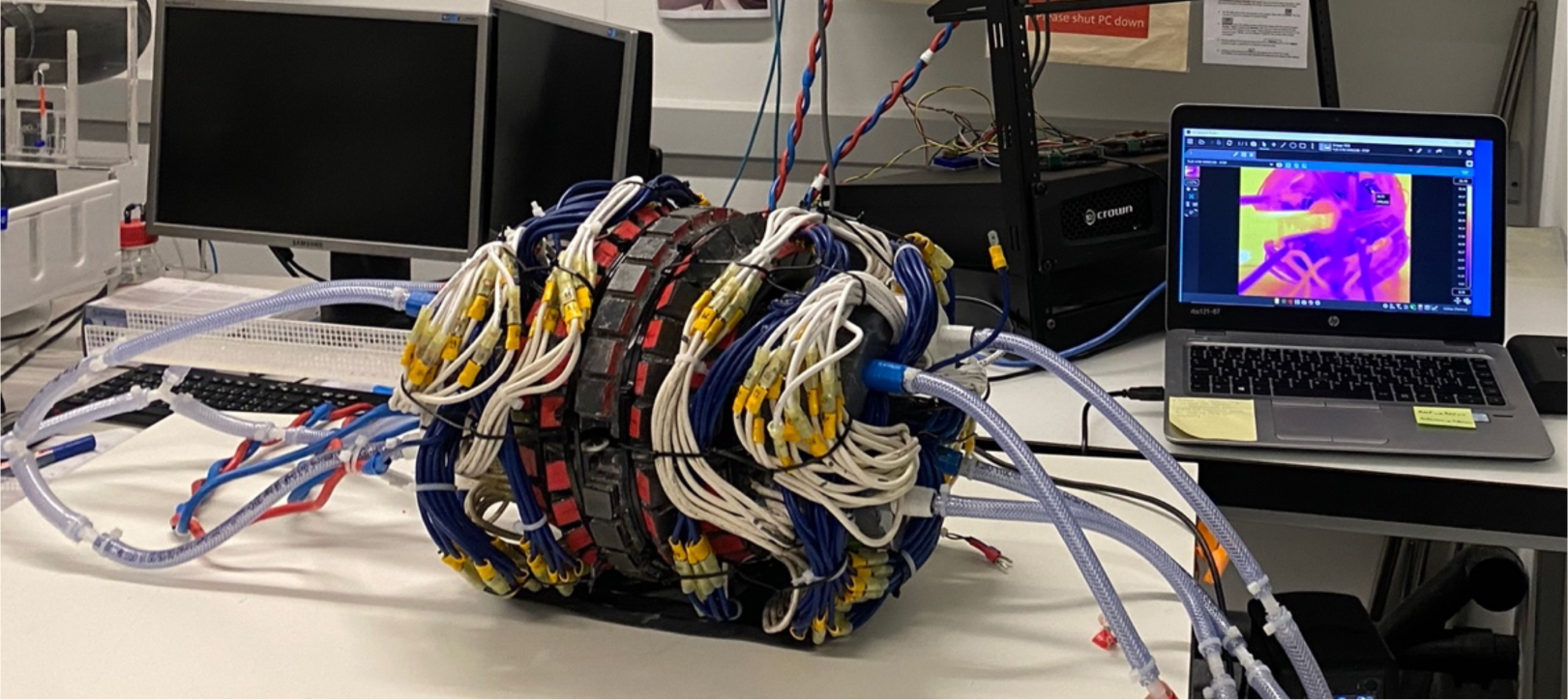
Photograph of the assembled prototype. An IR thermography readout monitoring the setup, and the class D audio amplifier powering the RMF coils are visible in the background. The Digilent devices that served as function generators are controlled with the computer and are sitting atop the audio amplifier.

